# BRCA1 promotes homologous recombination through separable DNA and RAD51 binding activities within its disordered region

**DOI:** 10.64898/2026.05.28.728454

**Authors:** Angela M. Jasper, Hoang H. Dinh, Wenjing Li, Qingming Fang, Cody M. Rogers, Sameer Salunkhe, Korilynn G. Kelly, Hardeep Kaur, Antoine Baudin, Xiaoping Xu, Tengyang Ni, Youngho Kwon, James M. Daley, Robert Hromas, Simon A. Gayther, Kate Lawrenson, Alexander V. Mazin, Sandeep Burma, Weixing Zhao, Patrick Sung, David S. Libich

**Author notes:** These authors contributed equally to this work (Angela M. Jasper, Hoang H. Dinh, Wenjing Li, Qingming Fang).

## Abstract

The breast and ovarian tumor suppressor BRCA1 heterodimerizes with BARD1 to promote DNA double-strand break repair by homologous recombination (HR) and to protect stressed DNA replication forks against nuclease attack. The large, intrinsically disordered central region of BRCA1 harbors binding sites for DNA and multiple repair factors, but its lack of stable structure has hindered mechanistic dissection of these activities. Using biochemical mapping and NMR spectroscopy, we delineate the DNA binding and RAD51 interaction interfaces within this region and construct separation-of-function mutants that selectively ablate each activity. Both DNA binding and RAD51 interaction are required for BRCA1-BARD1 to promote RAD51-mediated DNA strand invasion, and DNA binding also contributes to BLM-DNA2 end resection. These findings provide mechanistic insights into how individual ligand binding activities within BRCA1 contribute to genome maintenance.

## INTRODUCTION

*BRCA1* was identified as a familial breast cancer susceptibility gene more than three decades ago^1,2^, yet the mechanistic basis of BRCA1 protein function remains poorly understood. Pathogenic mutations in *BRCA1* predispose women to ovarian cancer and increase the risk of pancreatic and prostate cancers^3–5^. In addition, oncogenic somatic mutations occur across various cancer types^6–8^. At the cellular level, BRCA1 protein mutations cause hypersensitivity to DNA damaging agents and a pronounced deficiency in double-strand break (DSB) repair via HR^9,10^. BRCA1 is also involved in the protection of stressed DNA replication forks from nuclease attack^11–13^ and resolution of R-loops via the RNA-DNA helicase SETX^14,15^. Despite broad disease relevance and intensive interests, the mechanistic details that underpin BRCA1 roles in nuclear processes remain largely obscure.

BRCA1 functions in various biological processes as an obligate heterodimer with BARD1^16–18^, which is itself a suppressor of breast and other cancer types^19–21^. The BRCA1-BARD1 complex binds DNA and interacts physically with a variety of keystone repair factors, most notably CtIP, the RECQ family helicases BLM and WRN, the exonuclease EXO1, the recombinase RAD51, and with the tumor suppressor PALB2^17,18,22–24^. Through these interactions with nucleic acid and protein ligands, BRCA1-BARD1 exerts a broad, positive influence on multiple HR steps. In particular, BRCA1-BARD1 upregulates the activity of EXO1, BLM, and WRN to accelerate nucleolytic processing of DSBs, yielding a long 3’ ssDNA tail for assembly of RAD51-ssDNA nucleoprotein filaments capable of DNA homology search and strand invasion^18,25^. We have previously shown that BRCA1-BARD1 enhances the ability of RAD51-ssDNA nucleoprotein filaments to conduct DNA homology search and strand exchange^17^. Others have presented cellular evidence that BRCA1-BARD1 functions with BRCA2 and PALB2 to promote RAD51-ssDNA nucleoprotein filament assembly^23,26–29^.

Despite more than three decades of research, the importance of BRCA1-ligand interactions in HR and genome maintenance remains poorly understood. Defining the interfaces in BRCA1 that mediate nucleic acid binding and association with partner HR factors is essential for elucidating its roles in genome repair and maintenance. Progress has been limited, however, because of the large size of the BRCA1 protein (1,863 amino acid residues) and its mostly disordered nature. Here, we employ a combination of biochemical and high-resolution NMR analysis to identify key residues that constitute the DNA binding and RAD51 interaction regions within a central, disordered region of BRCA1. Being guided by the results of these analyses, we have constructed separation-of-function (SOF) mutants that specifically ablate each ligand binding activity and tested these SOF mutants to define how BRCA1 interactions with protein and nucleic acid ligands contribute to its genome maintenance functions.

## RESULTS

### BRCA1 disordered region harbors DNA and RAD51 binding sites

The protein segment encoded by exon 11 accounts for more than 60% of BRCA1 and harbors the DNA binding and RAD51 interaction regions^3,22,24^ (Figure 1A). Disorder prediction using IUPred^30^ indicates that exon 11 encodes a large intrinsically disordered region (IDR) (Figure S1A). The structural plasticity of IDRs facilitates interactions with biomolecular partners through short stretches of 8–12 amino acids, known as short linear interaction motifs (SLiMs)^31^. The conformational flexibility of IDRs supports their function as molecular scaffolds, allowing SLiMs to adopt orientations crucial for target recognition. This disordered nature, however, poses major challenges in delineating functional significance of the ligand interaction interfaces of BRCA1.

**Figure 1.**
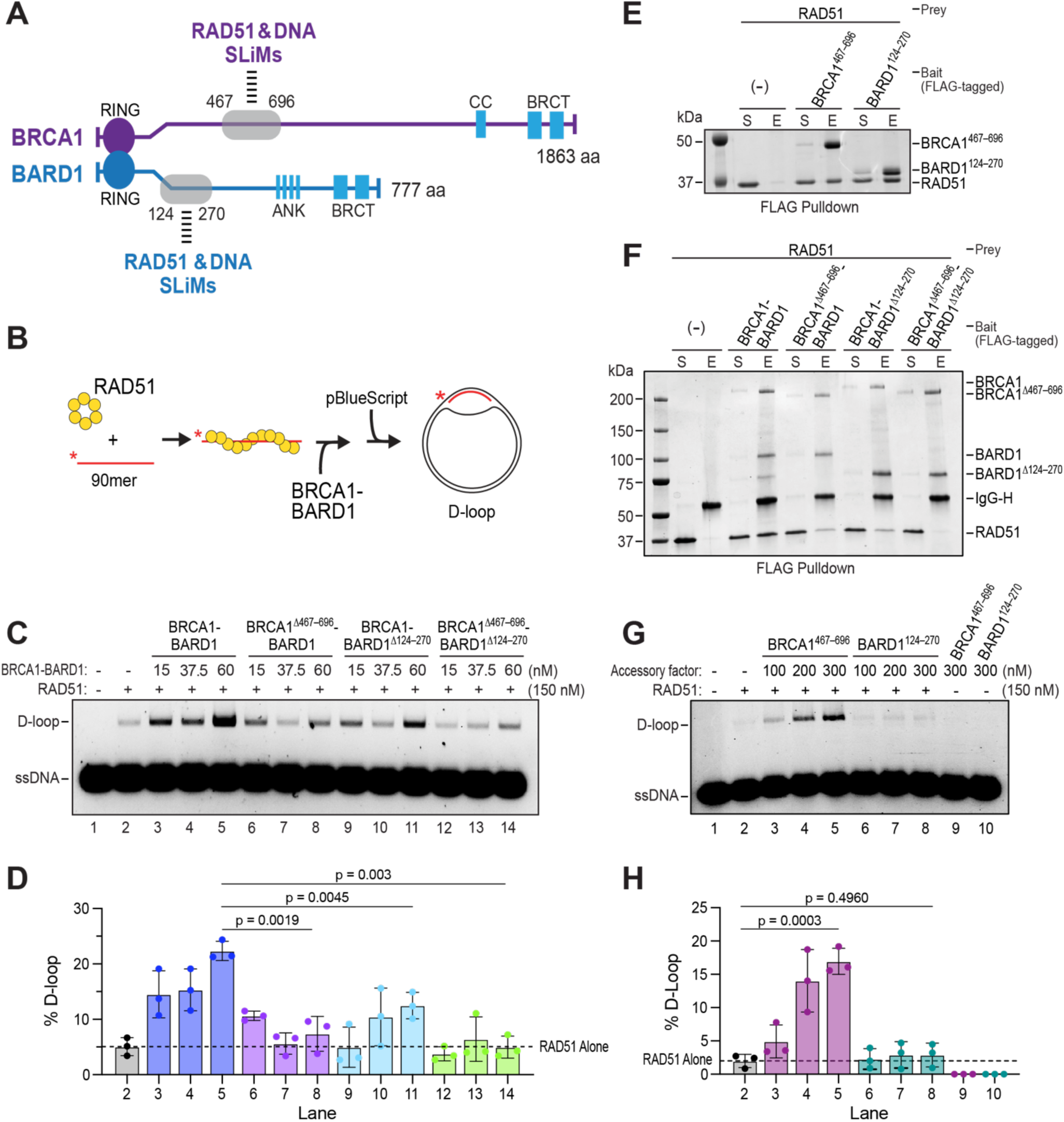
Roles of the BRCA1^467–696^ region in DNA binding, RAD51 interaction, and stimulation of D-loop formation. (A) Schematic of the domain structure of the BRCA1-BARD1 complex and its functional regions highlighting the regions (BRCA1^467-696^, BARD1^124-270^) encompassing DNA and RAD51 binding short linear interaction motifs (SLiMs). (B) Schematic of the D-loop assay. (C) RAD51-mediated D-loop reactions performed with BRCA1-BARD1 or the indicated deletion mutants. (D) Quantification of D-loop data from C; n = 3; mean ± SD (standard deviation). (E) Testing of FLAG-tagged BRCA1^467–696^ and BARD1^124-270^ for RAD51 interaction by affinity pulldown. Supernatant (S) containing unbound proteins and SDS eluate (E) were analyzed by SDS-PAGE and Coomassie blue staining. (F) Testing of FLAG-tagged BRCA1-BARD1 and the indicated deletion mutants for RAD51 interaction by affinity pulldown. Analysis was performed as in E. (G) Testing of BRCA1^467–696^ and BARD1^124–270^ for stimulation of RAD51-mediated D-loop formation. (H) Quantification of results from G; n = 3; mean ± SD.

Previous work established that BRCA1 binds DNA^10,17,18,24^, and that the N-terminal RING, C-terminal coiled-coil (CC), and tandem BRCT regions have minimal DNA binding activity^24^. To identify the DNA binding region of BRCA1, we expressed and purified protein fragments derived from exon 11 (Figures S1A-B) and tested them for DNA binding using the electrophoretic mobility shift assay (EMSA). Three overlapping fragments, BRCA1^360–696^, BRCA1^467–696^, and BRCA1^500–894^, exhibited DNA binding activity, with the first two showing higher affinity for DNA (Figure S1C). This analysis thus revealed that BRCA1^467–696^ encompasses the major DNA binding region of BRCA1.

BRCA1 physically interacts with RAD51^17,22^. Testing of the BRCA1 fragment library revealed that all fragments containing BRCA1^467–696^ interact with RAD51 (Figure S1D). Turbonuclease treatment or ethidium bromide inclusion confirmed that this interaction is not bridged by nucleic acids (Figure S1E). We previously showed that the BARD1 region encompassing residues 124–270 harbors DNA binding and RAD51 interaction activities^17^. Based on the biochemical mapping information, we constructed three deletion mutants within the context of full-length BRCA1-BARD1, namely BRCA1^Δ467–696^ (residues 467–696 deleted), BARD1^Δ124-270^ (residues 124–270 deleted), and the double deletion mutant (Figure S1F). As expected, all three BRCA1-BARD1 mutants are impaired for DNA binding activity (Figures S1G-H)^18^ and RAD51 interaction (see next section), with the most severe defects observed with the BRCA1^Δ467–696^-BARD1^Δ124-270^ double mutant.

### Functional relevance of BRCA1 DNA binding and RAD51 interaction activities

The BRCA1-BARD1 internal deletion mutants, the double deletion in particular, are impaired in their ability to stimulate the DNA end resection machinery^18^. Beyond its role in resection, BRCA1-BARD1 directly promotes RAD51-mediated D-loop formation^17^. We therefore tested the impact of the three BRCA1-BARD1 deletion mutations on this activity using a D-loop formation assay (Figure 1B). All three BRCA1-BARD1 mutants were impaired for RAD51 enhancement, with the greatest deficiency associated with the BRCA1^Δ467–696^-BARD1^Δ124-270^ double mutant (Figures 1C-D).

Since the deleted regions in BRCA1^Δ467–696^ and BARD1^Δ124-270^ also contain RAD51 interaction sites (Figure 1E) and BRCA1-BARD1 enhancement of D-loop formation is highly specific for human RAD51^17^, we tested the deletion mutants for RAD51 interaction to determine whether the observed defect in D-loop stimulation was solely attributable to loss of DNA binding. Consistent with the mapping data, BRCA1^Δ467–696^-BARD1 and BRCA1-BARD1^Δ124–270^ showed diminished association with RAD51, while the BRCA1^Δ467–696^-BARD1^Δ124–270^ double mutant was nearly devoid of RAD51 interaction capability (Figure 1F). These results indicate that the impaired ability of the BRCA1-BARD1 internal deletion mutants to enhance RAD51-mediated D-loop formation reflects loss of both DNA binding and RAD51 interaction.

In DNA end resection mediated by EXO1 and BLM-DNA2, BRCA1^467–696^ and BARD1^124–270^ function as standalone modules capable of stimulating resection activity^18^. We therefore asked whether these same fragments could stimulate RAD51-mediated D-loop formation. BRCA1^467–696^ elevated D-loop formation efficiently but only at concentrations higher than required for it to intact with the BRCA1-BARD1 complex, while BARD1^124–270^ showed minimal activity even at the highest concentrations tested (Figures 1G-H). Together, these results support the premise that the BRCA1^467–696^ region is required for BRCA1-BARD1 to enhance RAD51-mediated DNA strand invasion.

### Identification of the BRCA1 DNA binding interface and construction of mutants

To delineate the contribution of BRCA1 DNA binding to DNA damage repair and replication fork preservation, we designed SOF mutations that impair DNA binding without affecting RAD51 interaction. Sequence alignment of several BRCA1 orthologs identified conserved positively charged residues that could mediate electrostatic interactions with the negatively charged phosphodiester DNA backbone (Figure 2A). We selected six conserved lysine residues as mutagenesis targets and generated a DNA binding-deficient mutant (BRCA1^DM1^) by mutating K581, K583, K651, K652, K653, and K654 to glutamic acid (Figure 2A).

**Figure 2.**
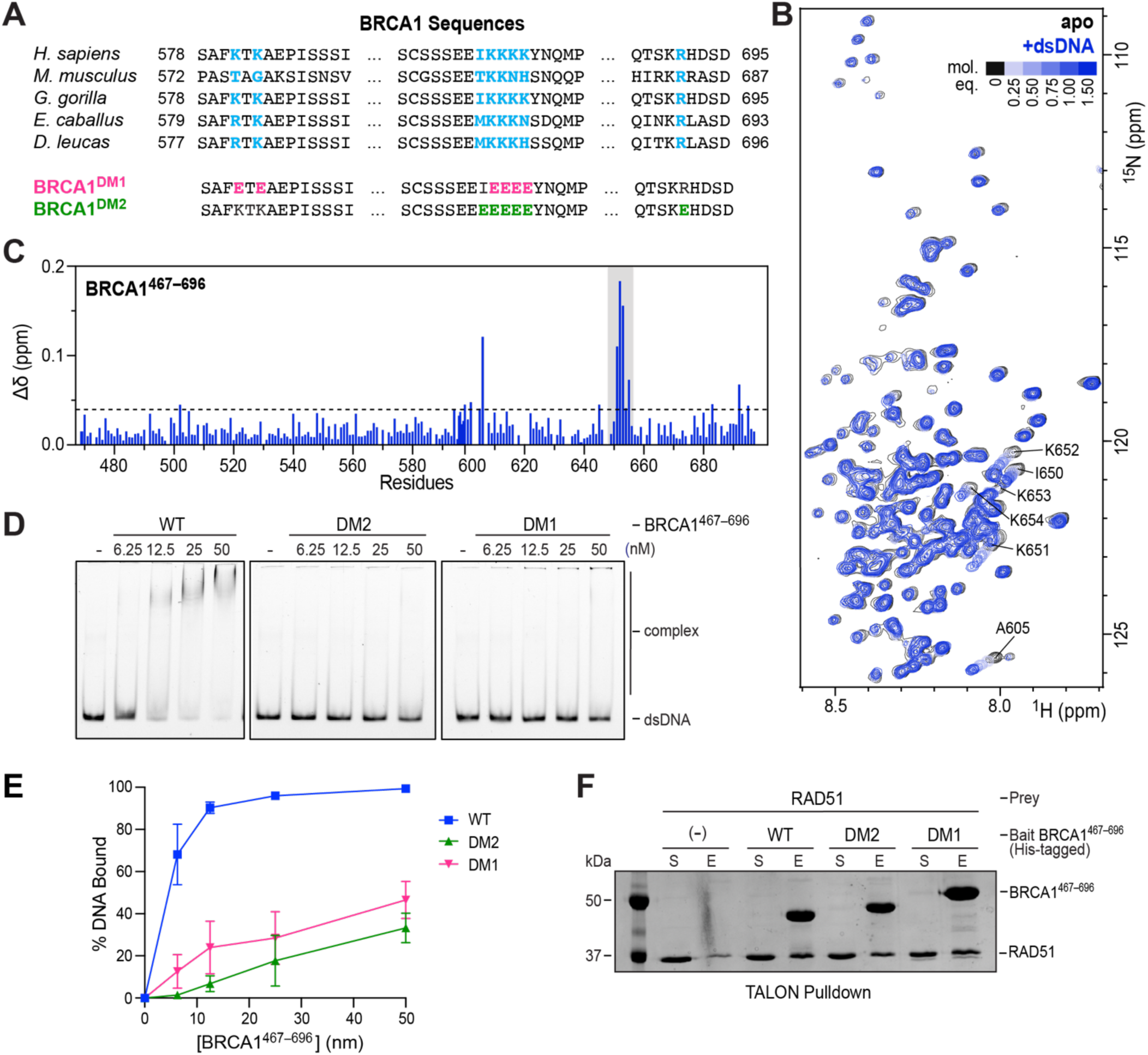
Delineation of the DNA binding interface in BRCA1 and mutant construction. (A) Alignment of human BRCA1 with orthologs, with conserved positively-charged residues highlighted in light blue. (B) Overlay of ^1^H,^15^N-HSQC spectra of ^15^N-BRCA1^467–696^ in the absence (black) or presence of increasing amounts of 20-mer duplex DNA (light blue to dark blue; 0.25 to 1.5 molar equivalents). Labeled peaks indicate residues with the most significant chemical shift perturbations. (C) Chemical shift perturbations (Δδ) within BRCA1^467–696^ induced by 1.5 molar equivalents of 20-mer duplex DNA. The dashed line represents the mean plus one standard deviation of Δδ for all peaks; shifts above this threshold are considered significant. The region mutated to yield the DM1 and DM2 mutants is highlighted in gray. (D) EMSA testing BRCA1^467–696^ wild-type, DM1, and DM2 with Cy5-labeled 80-mer dsDNA (1 nM). (E) Quantification of results from E; n = 3; mean ± SEM (standard error of the mean). (F) Affinity pulldown testing His-tagged BRCA1^467–696^ DM1 and DM2 for RAD51 interaction. Supernatant (S) and SDS eluate (E) were analyzed by SDS-PAGE and Coomassie blue staining.

We also utilized NMR, well suited to defining ligand interaction interfaces in disordered regions of target proteins^32,33^, to map the DNA binding interface within BRCA1^467–696^. The ^1^H,^15^N-HSQC spectrum of ^15^N-BRCA1^467–696^ revealed narrow ^1^H chemical shift dispersion, consistent with this fragment being an IDR (Figure 2B). Backbone resonance assignment was completed for 96% of residues (excluding prolines), and secondary structure propensity (SSP) calculations from ^13^C chemical shift data indicated an absence of stable secondary structure elements (Figure S2). Average ^15^N relaxation rates (*R*_1_ ∼ 1.6 s^-1^, *R*_2_ ∼ 3.4 s^-1^) and near zero ^1^H-^15^N heteronuclear NOE (hetNOE) values confirmed the fragment is largely disordered (Figure S2). To identify residues involved in DNA binding, we titrated duplex DNA into ^15^N-BRCA1^467–696^. Chemical shift perturbations mapped to a contiguous stretch of four consecutive lysine residues, K651–K654, consistent with a specific DNA interaction surface (Figures 2B-C). Notably, K651–K654 overlap with four of the six residues targeted in the DM1 mutant. To construct a second, NMR-guided mutant, we mutated K651–K654 along with I650 and R691, which also showed significant chemical shift perturbations, to glutamic acid (BRCA1^DM2^; Figure 2A).

By EMSA, we showed that BRCA1^467–696^ harboring either the DM1 or DM2 mutation is strongly impaired for dsDNA binding compared to the wild-type counterpart (Figures 2D-E). This impairment extended to ssDNA and fork DNA substrates. (Figures S3A-E). Because BRCA1^467–696^ also harbors the RAD51 interaction region (Figure 1E), we used affinity pulldown to test the DM1 and DM2 mutants for physical interaction with RAD51 (Figure 2F). Importantly, both mutants retain full RAD51 interaction capability, attesting to the SOF nature of these DNA binding mutations.

### Delineation of the RAD51 interaction interface and construction of mutants

Affinity pulldown results presented above indicate that BRCA1^467–696^ interacts with RAD51 (Figure 1E). By NMR analysis, we identified amino acid residues that constitute the RAD51 interaction interface. In the presence of two molar equivalents of RAD51, uniformly reduced peak intensities were observed across the entire sequence of BRCA1^467–696^, with an average I/I₀ value of 0.49. However, more pronounced line broadening was observed in two regions, residues 467–520 and 600–635 (Figures 3A-B). This differential broadening likely reflects direct contact between these BRCA1 regions and RAD51, arising from lifetime line broadening, conformational exchange, or a combination of both. We then conducted sequence alignment to identify conserved residues within these regions (Figure 3C). Conserved residues showing the greatest line broadening were selected as likely mediators of RAD51 interaction and targeted for mutagenesis. Accordingly, we constructed two mutants to ablate RAD51 interaction: RM1 (K501A, K503A, R504A, K505A, R506A, R507A) and RM2 (V626E, V627E, S628A, R629A, N630A) (Figure 3C). Affinity pulldown confirmed that both mutants are strongly impaired for RAD51 interaction (Figure 3D). Both BRCA1^RM1^ and BRCA1^RM2^ bind dsDNA with affinity comparable to the wild-type counterpart (Figures 3E-F), and this was also observed with ssDNA substrates (Figures S3F-H), thus demonstrating SOF between the DNA and RAD51 mutants.

**Figure 3.**
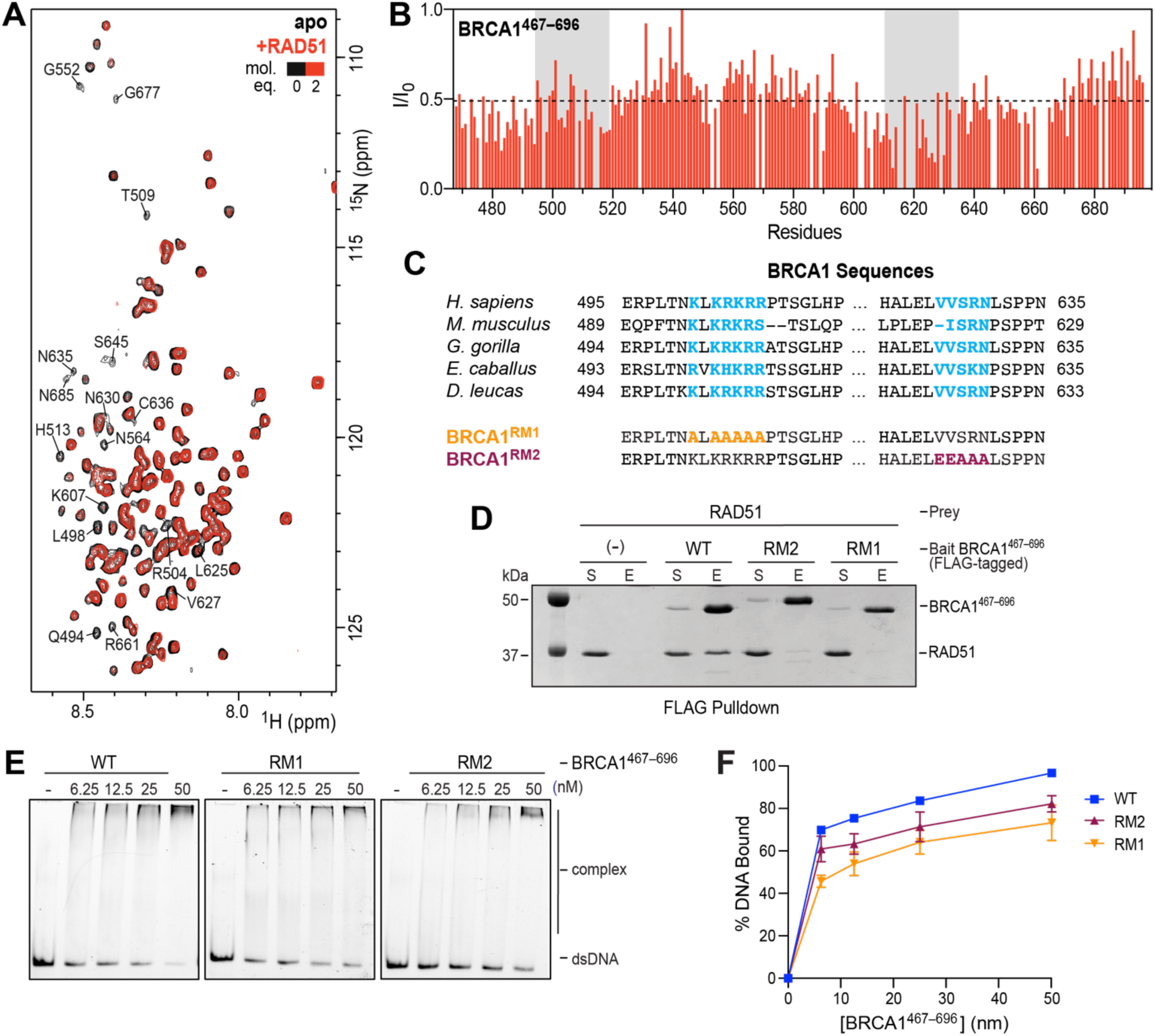
Delineation of the RAD51 interaction interface in BRCA1 and mutant construction. (A) Overlay of ^1^H,^15^N-HSQC spectra of ^15^N-BRCA1^467–696^ in the absence (black) or presence of 2 molar equivalents of RAD51 (red). Labeled peaks indicate residues with notable loss of peak intensity upon addition of RAD51. (B) Change in peak intensity (I/I₀) of residues in BRCA1^467–696^ induced by 2 molar equivalents of RAD51. The dashed line represents the mean minus one standard deviation of I/I₀; intensity ratios below this threshold are considered significant. Gray shading indicates the two regions with the greatest line broadening targeted for mutagenesis. (C) Alignment of human BRCA1 with orthologs highlighting conserved residues within the RAD51 interaction regions identified by NMR. Evolutionarily conserved residues are highlighted in blue; residues mutated in RM1 and RM2 are shown in orange and maroon, respectively. (D) Affinity pulldown testing FLAG-tagged BRCA1^467–696^ wild-type, RM1, and RM2 for RAD51 interaction. Supernatant (S) and SDS eluate (E) were analyzed by SDS-PAGE and Coomassie blue staining. (E) EMSA testing of BRCA1^467–696^ wild-type, RM1, and RM2 with Cy5-labeled 80-mer dsDNA (1 nM). (F) Quantification of results from E; n = 3; mean ± SEM.

### Role of BRCA1 DNA binding and RAD51 interaction in DNA strand invasion

To assess the contribution of BRCA1 DNA binding to promotion of DNA strand invasion, we generated recombinant BRCA1^DM1^-BARD1 and BRCA1^DM2^-BARD1 complexes (Figure S4A) using our established insect cell expression system. Mass photometry confirmed that these complexes were monodispersed with the expected 1:1 stoichiometry (Figure S4B). Testing in the D-loop assay revealed that BRCA1^DM1^-BARD1 and BRCA1^DM2^-BARD1 are both impaired for stimulation of strand invasion with D-loop yields reduced to levels comparable to RAD51 alone (Figures 4A-B), demonstrating that DNA binding by BRCA1 is required for BRCA1-BARD1 to promote DNA strand invasion.

**Figure 4.**
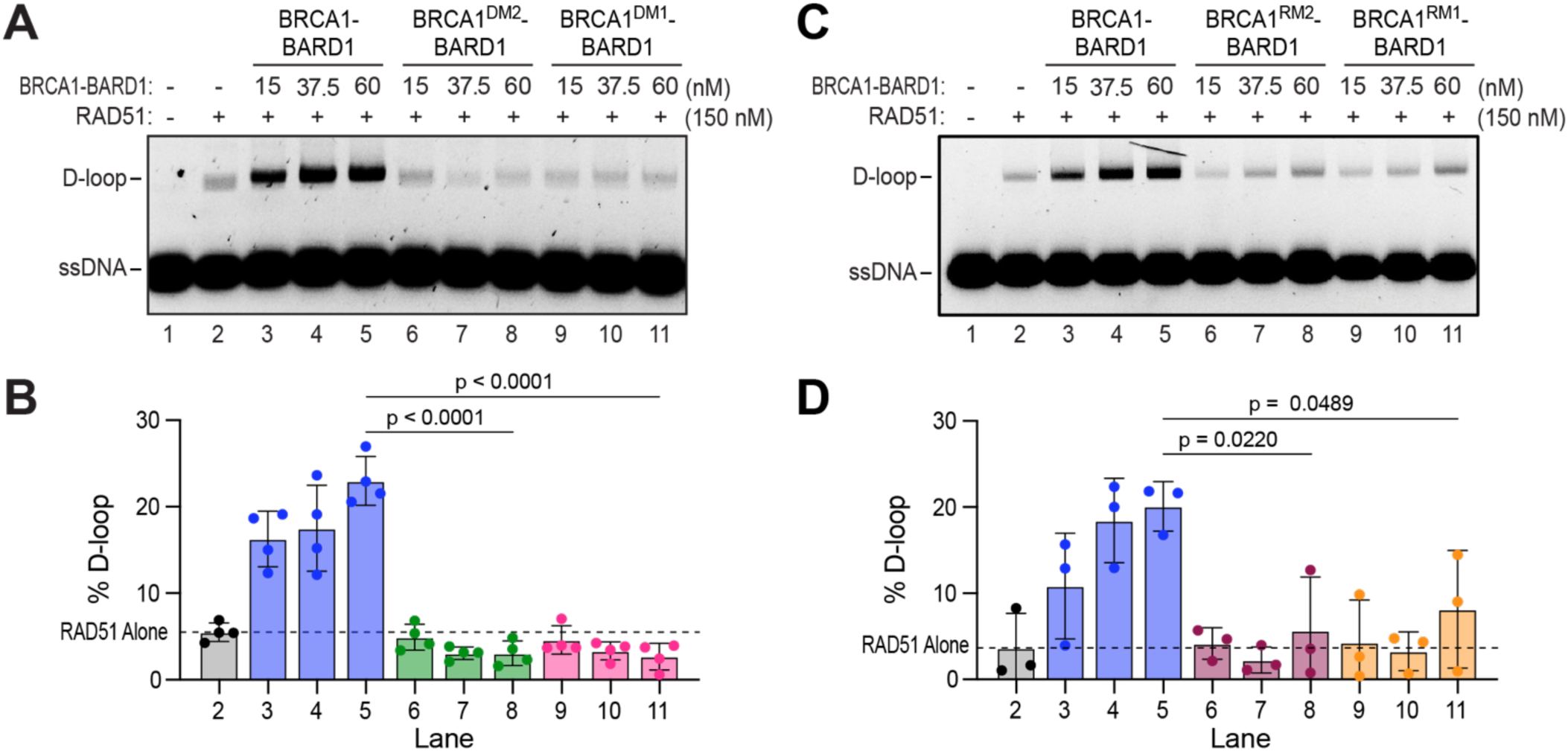
Role of BRCA1 DNA binding and RAD51 interaction in DNA strand invasion. (A) Testing of BRCA1^DM1^-BARD1 and BRCA1^DM2^-BARD1 tested alongside wild-type BRCA1-BARD1 for stimulation of RAD51-mediated D-loop formation. (B) Quantification of results from A; n = 3; mean ± SD. (C) Testing BRCA1^RM1^-BARD1 and BRCA1^RM2^-BARD1 tested alongside wild-type BRCA1-BARD1 for stimulation of RAD51-mediated D-loop formation. (D) Quantification of results from C; n = 3; mean ± SD. See Figure 1B for D-loop assay schematic.

To determine the contribution of RAD51 interaction by BRCA1 to D-loop formation, we generated full-length BRCA1^RM1^-BARD1 and BRCA1^RM2^-BARD1 complexes, again confirming their monodispersity and 1:1 stoichiometry by mass photometry (Figures S4C-D). The results showed that BRCA1^RM1^-BARD1 and BRCA1^RM2^-BARD1 are impaired for stimulation of D-loop formation, with D-loop yields markedly reduced relative to wild-type BRCA1-BARD1 (Figures 4C-D). Thus, loss of either DNA binding or RAD51 interaction by BRCA1 substantially diminishes the ability of BRCA1-BARD1 to stimulate D-loop formation, indicating that both activities are required for promotion of RAD51-mediated DNA strand invasion.

### Role of BRCA1 DNA binding in DNA end resection

We have previously shown that BRCA1-BARD1 stimulates long-range DNA end resection and that deletion of BRCA1 residues 467–696 modestly impairs stimulation of BLM-DNA2 end resection but has little impact on EXO1 activity^18,34^. We reasoned that this defect reflects reduced DNA binding by the BRCA1-BARD1 complex. To test this, we assessed BRCA1^DM1^-BARD1 and BRCA1^DM2^-BARD1 for stimulation of BLM activity. In the BLM helicase assay (Figure 5A), neither BRCA1^DM1^-BARD1 nor BRCA1^DM2^-BARD1 enhanced BLM helicase activity as robustly as wild-type BRCA1-BARD1 (Figures 5B-C). These DNA binding mutants were also similarly impaired for stimulation of BLM-DNA2-mediated end resection (Figures 5D-F). These deficiencies are intermediate in magnitude, consistent with our previously published analysis of the BRCA1^Δ467–696^-BARD1 mutant^18^. In contrast, BRCA1^RM1^-BARD1 and BRCA1^RM2^-BARD1 were just as proficient as wild-type BRCA1-BARD1 in stimulation of both BLM helicase activity (Figures 5B-C) and BLM-DNA2 end resection (Figures 5E-F). We have previously shown that the BRCA1^467–696^ region physically interacts with BLM. By affinity pulldown, both the DM and RM mutants retain BLM interaction comparable to wild type (Figures S5A-B), indicating that the deficiency observed with the DM mutants is attributable to loss of DNA binding rather than impaired BLM interaction.

**Figure 5.**
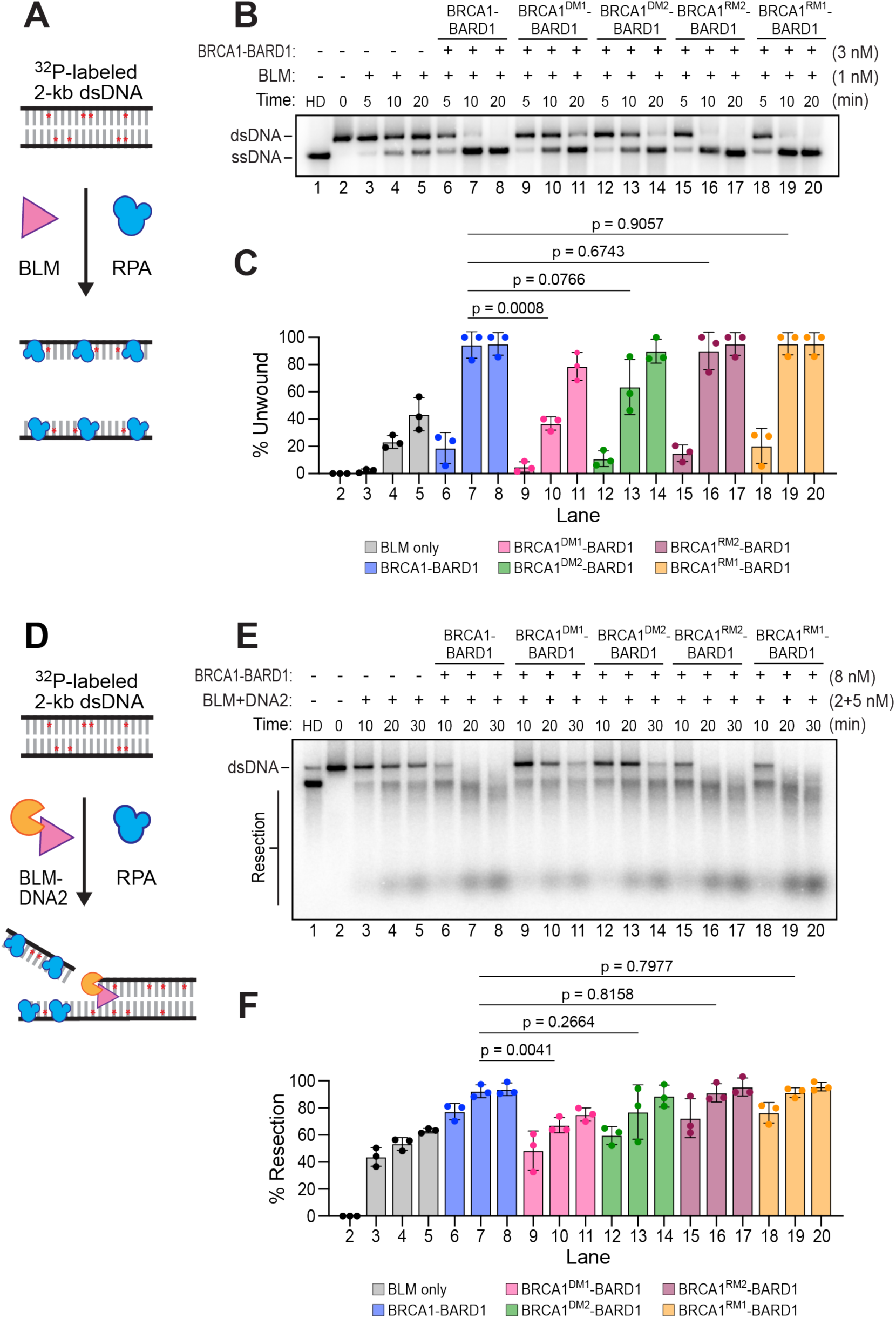
Role of BRCA1 DNA binding in BLM-mediated DNA unwinding and BLM-DNA2-mediated end resection. (A) Schematic of the DNA helicase assay. (B) Testing of BRCA1-BARD1 DNA binding (DM1, DM2) and RAD51 interaction (RM1, RM2) mutants tested alongside wild-type BRCA1-BARD1 for stimulation of DNA unwinding by BLM. (C) Quantification of the DNA unwinding results from B; n = 3; mean ± SD. (D) Schematic of the DNA end resection assay. (E) Testing of BRCA1-BARD1 DNA binding (DM1, DM2) and RAD51 interaction (RM1, RM2) mutants tested alongside wild-type BRCA1-BARD1 for stimulation of BLM-DNA2-mediated end resection. (F) Quantification of the DNA end resection results from E; n = 3; mean ± SD.

BRCA1^DM1^-BARD1 and BRCA1^DM2^-BARD1 are proficient in stimulation of EXO1-mediated end resection (Figures S5C-E), consistent with prior work showing that deletion of the BRCA1 DNA binding region elicits only a mild deficiency in EXO1 stimulation by BRCA1-BARD1^18,34^. However, the DM1 and DM2 mutations ablate the ability of the BRCA1^467–696^ fragment to enhance EXO1-mediated resection (Figures S5F-G), indicating that DNA binding by this region of BRCA1 does contribute to EXO1 stimulation, but its contribution within the context of the BRCA1-BARD1 complex is modest. These data, combined with our previous work^18,34^, support the conclusion that BARD1 DNA binding alone is sufficient for EXO1 upregulation by the BRCA1-BARD1 complex.

## DISCUSSION

BRCA1 acts at multiple stages of HR and protects stressed replication forks against nucleolytic attrition, but mechanistic dissection of these functions has been hampered by the large size and intrinsically disordered nature of BRCA1. Here, we combine biochemical mapping and NMR spectroscopy to delineate the DNA binding and RAD51 interaction interfaces within the BRCA1 disordered region and construct SOF mutants that specifically ablate each activity. Using these mutants, we demonstrate that both DNA binding and RAD51 interaction by BRCA1 are required for stimulation of RAD51-mediated DNA strand invasion, and that DNA binding also contributes to BLM-DNA2 end resection. (Figure 6).

**Figure 6.**
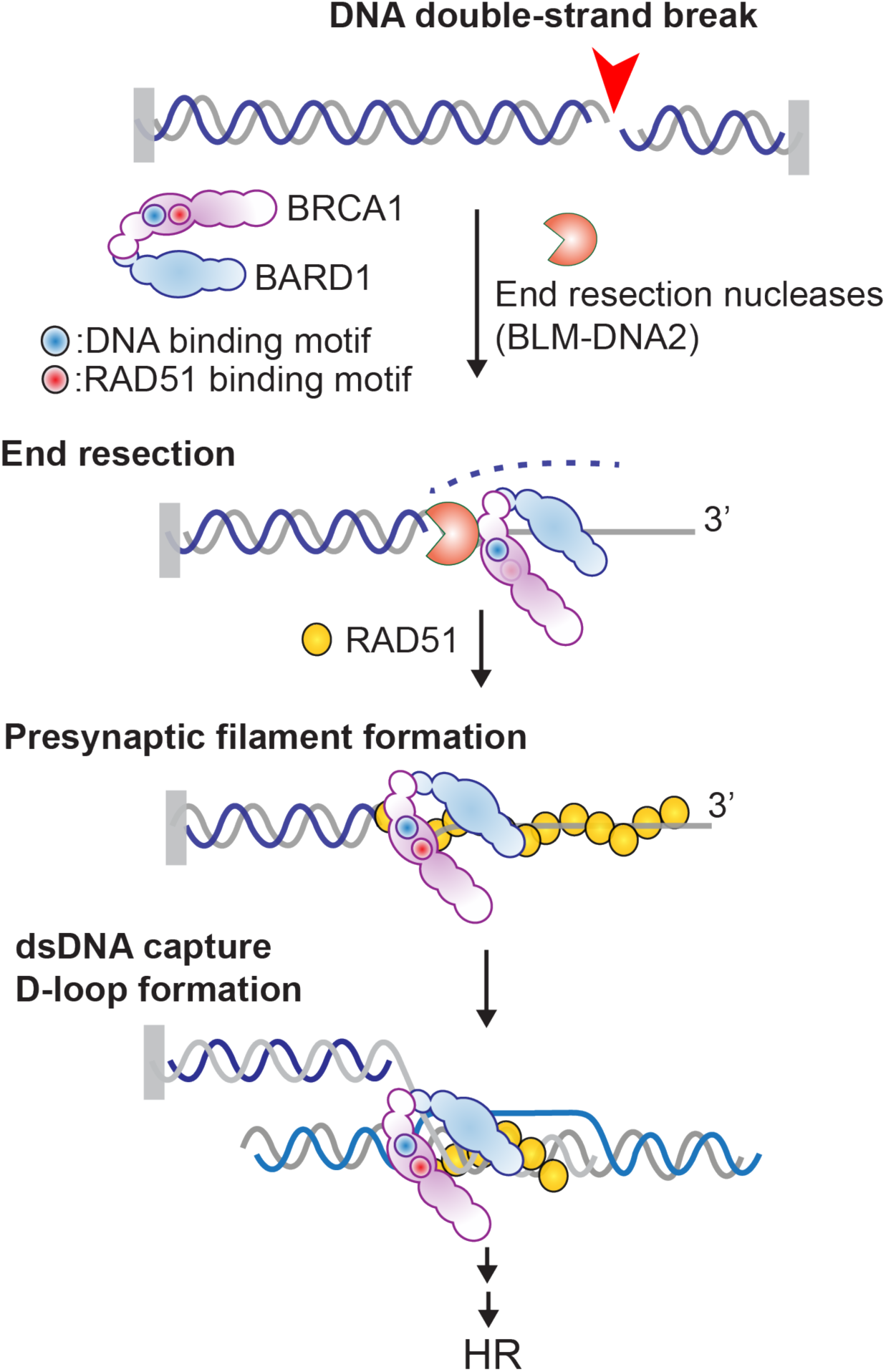
Model of BRCA1 DNA binding and RAD51 interaction attributes in genome maintenance. BRCA1 (purple) and its obligate binding partner BARD1 (blue) promote multiple genome maintenance processes including homologous recombination (HR) at DNA double-strand breaks. DNA binding by BRCA1 is required for stimulation of BLM-DNA2-mediated end resection and RAD51-mediated D-loop formation. RAD51 interaction by BRCA1 is required for stimulation of D-loop formation and protection of stalled replication forks.

The BRCA1^467–696^ region represents a multifunctional module capable of enhancing RAD51-mediated DNA strand invasion and, as we have previously shown, upregulating the DNA end resection machinery including BLM, WRN, and EXO1^18^. The intrinsically disordered nature of this region likely underpins this functional versatility. In the fly-casting model^35^, the extended conformation of an IDR increases the effective capture radius of the protein, allowing it to sample a larger volume of space and thereby enhancing the likelihood of encountering specific binding partners. Within this framework, multiple SLiMs^31^ distributed across the disordered region collectively form high-affinity interaction interfaces for individual ligands such as DNA and RAD51. Notably, both BRCA1 and BARD1 possess SLiM-based DNA binding and RAD51 interaction activities, raising the possibility that the two subunits function cooperatively within the heterodimer. If the SLiMs in BRCA1 and BARD1 have distinct affinities for these ligands, the heterodimer may be capable of discriminating between different substrate contexts, such as the mixed DNA structures encountered at replication forks and resected DSBs.

Our findings provide the requisite foundation to investigate other BRCA1-dependent processes in genome maintenance. The SOF mutants and the combined biochemical and NMR-based approach used in this study can be extended to other ligands that interact with this disordered region, including BLM, WRN, and RAD50^18,25,36^, enabling the design of mutations that ablate specific binding activities or interrogate interfaces where multiple ligands compete or cooperate. This approach can also be applied to other BRCA1-dependent pathways, including R-loop resolution and the mechanism by which BRCA1 overcomes the anti-HR activity of the 53BP1-Shieldin axis to promote HR over NHEJ^10,34,37^. Notably, many key DNA repair factors, including 53BP1 (>1,900 residues, >80% disordered), an agonist of BRCA1^38^, share the large, disordered, multivalent architecture that has made BRCA1 challenging to study. The approach developed here thus provides the framework for dissecting the functional contributions of individual interaction interfaces within 53BP1 and other regulatory factors that harbor large IDRs. Within the context of higher order protein complexes, these IDRs are likely to compete or cooperate for ligand binding, regulating protein function in the context of large DNA repair complexes. More broadly, defining the interaction interfaces of BRCA1-BARD1 and their contributions to DNA damage repair and replication fork preservation will help illuminate the functional consequences of cancer-associated mutations and inform the development of therapeutic strategies targeting repair-deficient tumors.

## MATERIALS AND METHODS

### Plasmids

The DNA coding for BRCA1^467–696^ fragments were synthesized and cloned into pET-3a or pE-SUMO vectors by Gene Universal. Mutations were introduced by QuikChange site-directed mutagenesis or commercially synthesized and cloned into the pET-3a expression vector by Gene Universal. RAD51 and RPA were expressed in bacteria using previously described vectors^18,39^. Baculoviruses for insect cell expression of BRCA1, BARD1, EXO1, BLM, and DNA2 were generated from vectors described previously^17,18,40^. A modified HA-BRCA1 mammalian cell expression vector^40^ containing an additional FLAG tag was used to generate stable cell lines. All constructs were verified by sequencing to confirm the absence of unintended mutations.

### Protein Expression and Purification

*BRCA1 fragments:* Plasmids of the expression of His-SUMO-BRCA1 and His-SUMO-BRCA1-FLAG fragments (wild-type or mutant) were transformed into Rosetta (DE3) *E. coli* cells. Cultures were grown in Luria-Bertani (LB) media at 37°C to an optical density of 0.6, induced with 0.5 mM isopropyl β-D-thiogalactopyranoside (IPTG), and grown for 16 hours at 16°C. Cells were harvested by centrifugation and stored at −80°C. All purification steps were carried out at 0-4°C. Cell pellets were resuspended and sonicated in 50 mL of T buffer (25 mM Tris-HCl pH 7.5, 10% glycerol, 0.5 mM EDTA, 0.05% IGEPAL CA-630 (Sigma-Aldrich), 2 mM DTT, and protease inhibitors (5 μg/mL each of aprotinin, chymostatin, leupeptin, and pepstatin)) supplemented with 500 mM KCl. The lysate was clarified by centrifugation at 100,000 x g for 45 min and incubated with 2 mL of Ni-NTA resin (Qiagen) for 1 hour. The resin was washed sequentially with 100 mL of T buffer containing 1 M KCl, 20 mM imidazole, 1 mM ATP, and 2 mM MgCl_2_, followed by 20 mL of T buffer with 300 mM KCl. Bound proteins were eluted by five sequential 10-min incubations with 2 mL of T buffer containing 300 mM KCl and 200 mM imidazole. Where indicated, the His-SUMO tag was cleaved by overnight incubation with Ulp1 protease at 4°C. Protein fragments were then fractionated on a 24 mL Superdex 200 column (GE) in T buffer with 300 mM KCl and concentrated using an Amicon device with a 10 kDa molecular weight cut-off.

### Isotope-enriched BRCA1^467–696^ fragment

His-SUMO-BRCA1^467–696^ in the pE-SUMO vector was transformed into BL21 Star (DE3) *E. coli* cells. Cultures were grown at 37°C in M9 media supplemented with appropriate isotopes (^15^NH_4_Cl, ^13^C-glucose) to an OD_600_ of 0.6, induced with 0.5 mM IPTG, and grown for an additional 4 hours at 37°C. Cells were harvested by centrifugation and stored at −80°C. All purification steps were carried out at 0-4°C. Cell pellets were resuspended in 50 mL of T buffer (25 mM Tris-HCl pH 7.5, 10% glycerol, 1 mM DTT, 0.5 mM EDTA, 0.05% IGEPAL CA-630) supplemented with Turbonuclease (Sigma-Aldrich, 500 units), protease inhibitors (5 μg/mL each of aprotinin, chymostatin, leupeptin, and pepstatin), 2 mM MgCl_2_, 20 mM imidazole, and 500 mM KCl. Cell lysate was prepared by sonication and clarified by centrifugation (100,000 x g, 45 min). Clarified lysate was incubated with 2 mL of Ni-NTA resin (Qiagen) for 1 hour. The resin was washed with 100 mL of T buffer supplemented with 2 mM MgCl_2_, 1 mM ATP, 20 mM imidazole, and 1 M KCl, followed by 20 mL of T buffer with 20 mM imidazole and 500 mM KCl. Bound protein was eluted five times with 2 mL of T buffer containing 200 mM imidazole and 500 mM KCl. Elution fractions were pooled and incubated overnight with Ulp1 to cleave the His_6_-SUMO tag. Pooled fractions were buffer exchanged into T buffer with 500 mM KCl using an Amicon 10K concentrator and incubated with 0.5 mL Ni-NTA resin for 30 min to remove cleaved tag. Flow-through was collected, buffer exchanged into Buffer N (20 mM KH_2_PO_4_ pH 6.5, 150 mM KCl, 0.5 mM DTT, 0.2 mM EDTA), concentrated to 0.5 mL, and fractionated on a Superdex Increase 75 10/300 column (Cytiva) with 24 mL of Buffer N. Peak fractions were pooled, concentrated to approximately 600 μL, snap frozen in liquid nitrogen, and stored at −80°C.

#### RAD51

His_6_-SUMO-RAD51 was expressed in Rosetta (DE3) *E. coli* cells transformed with pET-Duet-his-smt3-Rad51^39^. Cultures were grown at 37°C to an OD_600_ of 0.6, induced with 0.5 mM IPTG, and grown overnight at 16°C. Cells were harvested by centrifugation and stored at −80°C. All purification steps were carried out at 0-4°C. Cell pellets were resuspended in 50 mL of T buffer supplemented with Turbonuclease (Sigma-Aldrich, 500 units), protease inhibitors (5 μg/mL each of aprotinin, chymostatin, leupeptin, and pepstatin), 2 mM MgCl_2_, and 500 mM KCl. Cell lysate was prepared by sonication and clarified by centrifugation (100,000 x g, 45 min). The clarified lysate was subjected to ammonium sulfate precipitation (0.277 g/mL lysate) and the precipitate was harvested by centrifugation (18,000 x g, 30 min). The precipitate was resuspended in T buffer with 500 mM KCl and 20 mM imidazole and incubated with 2 mL of Ni-NTA resin overnight. The resin was washed sequentially with 30 mL each of T buffer containing 500 mM, 300 mM, and 150 mM KCl. The resin was then resuspended in 10 mL of T buffer with 150 mM KCl and incubated overnight with Ulp1 to cleave the His_6_-SUMO tag. The resin was loaded onto a gravity column and flow-through was collected and concentrated to 5 mL using an Amicon 10K concentrator. Protein was further fractionated on a 1 mL HiTrap Q HP column (Cytiva) using a salt gradient of 100–600 mM KCl. Peak fractions were pooled, concentrated, aliquoted, and stored at −80°C.

### Other recombinant proteins

BLM, DNA2, EXO1, and RPA were expressed and purified as previously described^41–43^.

### DNA Substrates

5’-Cy5-labeled and unlabeled oligonucleotides used in this study are listed in Table 1 and were purchased from Integrated DNA Technologies (IDT). Unlabeled oligonucleotides longer than 60 nucleotides were purified by denaturing polyacrylamide gel electrophoresis (PAGE). Oligonucleotides were resuspended in 95% formamide dye and electrophoresed on 10% denaturing TAE (40 mM Tris base pH 8.4, 20 mM acetic acid, 1 mM EDTA) polyacrylamide gels containing 7 M urea at 110 V, 60°C for 4 hours. Oligonucleotides were visualized under ultraviolet light and the prominent band was excised. DNA was extracted from the gel by electroelution at 30 V for 16 hours at 4°C. Purity was assessed by native gel electrophoresis and ethidium bromide staining. Duplex and fork DNA substrates were generated by mixing equimolar quantities of complementary oligonucleotides in annealing buffer (20 mM Tris-HCl pH 7.5, 4% glycerol, 0.1 mM EDTA, 4 mg/mL BSA, 10 mM DTT, and 10 mM MgCl_2_), heating to 95°C, and cooling at 0.1°C/min to 10°C in a thermocycler. Annealing efficiency was assessed by native gel electrophoresis. Substrates were further purified from unannealed olionuceotides by native PAGE and electroelution.

**Table 1:**
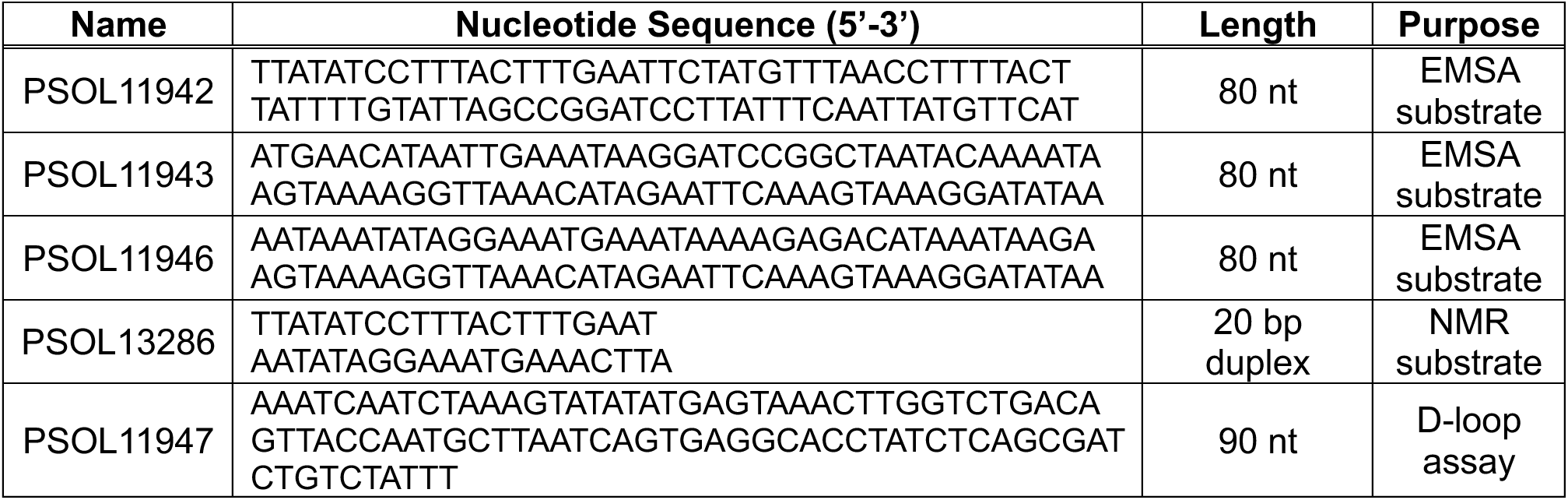
Sequences of oligonucleotides used to assess BRCA1 DNA binding.

### Electrophoretic mobility shift assays (EMSAs)

Duplex, fork, and bubble DNA substrates were assembled from oligonucleotides PSOL11942/PSOL11943, PSOL11942/PSOL11946, and PSOL11936/PSOL11937, respectively.

Oligonucleotide PSOL11942 or PSOL11936 was 5’-Cy5-labeled for visualization. These substrates and ssDNA (PSOL11942) at 1 nM were incubated with BRCA1-BARD1 (wild-type or indicated mutant) or BRCA1^467–696^ fragment (wild-type or indicated mutant) at 37°C for 5 min in 10 μL of buffer E (35 mM Tris-HCl pH 7.5, 1 mM MgCl_2_, 100 μg/mL BSA). Samples were mixed with loading dye (0.05% Orange G, 30% glycerol) and resolved by 5% native PAGE in TA buffer (30 mM Tris-acetate, pH 8.4) for BRCA1-BARD1 EMSAs or TB buffer (15 mM Tris-borate, pH 8.4) for BRCA1 fragment EMSAs.

### Affinity pulldowns

Affinity-tagged bait and prey proteins were incubated in 20 μL of buffer P (25 mM Tris-HCl pH 7.5, 1 mM DTT, 0.05% IGEPAL CA-630, 125 mM KCl) containing Turbonuclease (Sigma, 125 units) for 1 hour at 4°C. Reactions were then mixed with 5 μL of the indicated affinity resin for 30 min at 4°C. The resin was pelleted by centrifugation and 10 μL of supernatant was collected. The resin was washed three times with 50–150 μL of buffer P and eluted with 10 μL of 2x Laemmli buffer. The supernatant and eluate fractions were analyzed by SDS-PAGE with Coomassie blue staining or immunoblotting.

### D-loop assay

The D-loop assay was performed as previously described with minor modifications^17^. Cy5-labeled 90-mer oligonucleotide PSOL11947 (400 nM nucleotides) was incubated with RAD51 (150 nM) in buffer D (25 mM Tris-HCl, 1 mM DTT) containing 1 mM MgCl_2_ and 2 mM AMP-PNP at 37°C for 5 min. The indicated amount of BRCA1-BARD1 was then added followed by a 5 min incubation at 37°C. The D-loop reaction was initiated by the addition of pBluescript SK replicative form I DNA (12.3 μM base pairs) at a 1:1 molar ratio to the 90-mer and the completed reaction was incubated at 37°C for 10 min. Reactions were stopped and deproteinized by adding glycerol loading dye containing 1% SDS and 1 mg/mL proteinase K and incubated at 37°C for 10 min. Reactions were resolved in 1.1% agarose gels in TAE buffer (40 mM Tris base pH 8.4, 20 mM acetic acid, 1 mM EDTA) and imaged using a ChemiDoc Imaging System (Bio-Rad).

### Helicase and resection assays

DNA helicase and end resection assays were performed as previously described^18^. Briefly, reactions were carried out in buffer R (20 mM Na-HEPES pH 7.5, 2 mM ATP, 0.1 mM DTT, 100 μg/mL BSA, 0.05% Triton X-100, 2 mM MgCl_2_, and 100 mM KCl) with a 2-kilobase pair (2-kb) dsDNA substrate (0.5 nM ends) generated by PCR with internally labeled ^32^P-dCTP (Revvity)^43^. Reactions were incubated at 37°C for the indicated time, then stopped and deproteinized by addition of glycerol loading dye containing 1% SDS and 1 mg/mL proteinase K for 10 min at 37°C. Reactions were resolved on 0.9% agarose gels in TAE buffer (40 mM Tris base pH 8.4, 20 mM acetic acid, 1 mM EDTA), dried under vacuum, and analyzed by phosphorimaging (Amersham Typhoon).

### NMR spectroscopy

All experiments were recorded at 25°C on a Bruker Avance NEO spectrometer operating at a proton Larmor frequency of 700.13 MHz. All ^1^H,^15^N-HSQC spectra were acquired using 64* x 1024* complex data points in the indirect (^15^N) and direct (^1^H) dimensions, corresponding to acquisition times of 41.0 and 112.6 ms, respectively. Spectra were apodized with a sine bell function and zero-filled to twice the number of acquired points. Data were processed with Topspin 4.4.0 (Bruker) or NMRPipe^44^ and analyzed with CCPNMR Analysis 2.5.2^45^.

^15^N *R*_1_ and *R*_2_ relaxation rates were determined from *T*_1_ and *T*_1_ρ experiments recorded on a 125 μM sample of BRCA1^467–696^ in Buffer N. The ^15^N *T*_1_ experiment consisted of eight interleaved spectra with relaxation delays of 40, 80, 200, 280, 300, 400, 600, and 800 ms. The *T*_1_ρ experiment was recorded using a B_1_ field of 1400 Hz with eight interleaved spectra and relaxation delays of 1, 21, 31, 41, 61, 81, 121, and 161 ms. ^15^N *R*_2_ rates were calculated using the following equation:

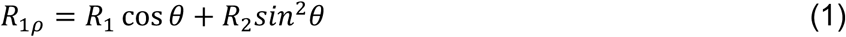

where θ = arctan(ω_1_/Ω), ω_1_ is the B_1_ field strength (1400 Hz) and Ω is the offset from the spinlock carrier frequency^46^. ^1^H-^15^N heteronuclear NOE experiments were recorded on the same sample and consisted of two interleaved experiments, with and without proton saturation, using a recycle delay of 4 s.

For DNA binding experiments, aliquots of unlabeled DNA (ssDNA, PSOL13287; dsDNA, PSOL13286; hairpin fork, PSOL13307) were titrated into 50 μM ^15^N-BRCA1^467–696^ in 500 μL of Buffer N in a 5 mm NMR tube (Wilmad) at BRCA1:dsDNA molar ratios of 4:1, 2:1, 3:4, 1:1, and 2:3. Chemical shift perturbations (CSPs) were calculated by weighting the ^1^H and ^15^N chemical shifts according to their gyromagnetic ratios^47^:

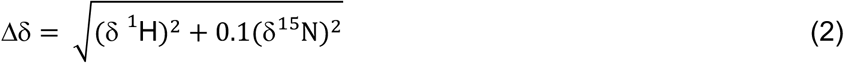

CSPs were considered significant when Δδ exceeded the sum of the mean and one standard deviation of Δδ for all residues.

For RAD51 titration, aliquots of 300 μM unlabeled RAD51 were titrated into 50 μM ^15^N-BRCA1^467–696^ in 500 μL of Buffer N in a 5 mm NMR tube (Wilmad) at BRCA1:RAD51 molar ratios of 1:1 and 1:2. Line broadening was assessed by calculating the change in peak intensity (I/I₀) between apo BRCA1^467–696^ and the highest RAD51:BRCA1 ratio recorded^48^.

### Mass photometry

Mass photometry was performed using a Refeyn TwoMP instrument. Samples were flash diluted to 20 nM in 1x phosphate-buffered saline (PBS). Data were acquired for 1 min using AcquireMP software (Refeyn) and mass distributions were analyzed with DiscoverMP software (Refeyn) v1.45.0. Ratiometric counts were converted to molecular weight (kDa) using a standard curve generated from ovalbumin (43 kDa), conalbumin (75 kDa), aldolase (158 kDa), and thyroglobulin (669 kDa)^18^.

## AUTHOR CONTRIBUTIONS

Conceptualization, A.M.J., H.H.D., C.M.R., H.K., P.S., and D.S.L.; Methodology, A.M.J., H.H.D., W.L., C.M.R., H.K., A.B., Y.K., W.Z., P.S., and D.S.L.; Investigation, A.M.J., H.H.D., W.L., Q.F., C.M.R., K.G.K., H.K., A.B., X.X., S.S., and T.N.; Writing – original draft, A.M.J. and P.S.; Writing – review & editing, A.M.J., H.H.D., W.L., Q.F., C.M.R., Y.K., P.S., and D.S.L; Funding acquisition, Y.K., S.B., A.V.M., P.S., W.Z., D.S.L.; Resources, A.M.J., H.H.D., W.L., C.M.R., H.K., X.X., T.N., J.M.D., R.H., S.G., K.L., and W.Z.; Supervision, R.H., S.G., K.L., A.V.M., S.B., W.Z., P.S., and D.S.L.

## ACKNOWLEDGMENTS

This study was supported by National Institutes of Health grants: R01 GM140127, R35 GM163922 (D.S.L.), R35 CA241801, R01 ES007061, R01 CA168635 (P.S.), P01 CA275717 (Y.K., P.S., S.B., D.S.L., A.V.M., W.Z., R.H.), R01 GM141091 (W.Z.), R01 CA246807 (S.B.), R01 CA237286, R01 GM136717 (A.V.M.), R01 CA251555, R01 CA259058, R01 CA251722 (S.A.G.), R01 CA13942 (R.H.), R50 CA265315 (Y.K.), F30 CA278370, T32 GM145432 (A.M.J.); American Cancer Society grants: RSG-22-721675-01-DMC, DBG-25-1292405-01-DMC (W.Z.), PF-22-034-01-DMC (C.M.R.); Cancer Prevention and Research Institute of Texas (CPRIT) grants RR210023 (A.V.M.), RR230078 (S.A.G.); Congressionally Directed Medical Research Programs: BC191160 (A.V.M), OC220326 (K.L, S.A.G.), HT9425-23-1-3262, HT9425-24-1-0716 (S.A.G.); The Welch Foundation AQ-2001-20190330 (D.S.L.). P.S. is the holder of the Robert A. Welch Distinguished Chair in Chemistry (AQ-0012). A.V.M. is a holder of the Joe R. and Teresa Lozano Long Chair in Cancer Research. H.H.D. and A.B. thank the Greehey Family Foundation and the Center for Biological Neuroscience, respectively, for fellowship support. The authors thank Dr. Timsi Rao for initial work on BRCA1 DNA binding, and Dr. Kristin Cano for technical assistance with NMR experiments. This work is based upon research conducted in the Structural Biology Core Facilities, a part of the Institutional Research Cores at the University of Texas Health Science Center at San Antonio supported by the Office of the Vice President for Research and the Mays Cancer Center Drug Discovery and Structural Biology Shared Re-source (NIH P30 CA054174)

## Data Availability

Resonance assignments for BRCA1^467–696^ are available in the Biological Magnetic Resonance Data Bank under accession number 53585. Further information and requests for resources and reagents should be directed to and will be fulfilled by the lead contact, David S. Libich.

## Abbreviations

