## Supplemental Material for "BRCA1 promotes homologous recombination through separable DNA and RAD51 binding activities within its disordered region"

### SUPPLEMENTARY MATERIAL

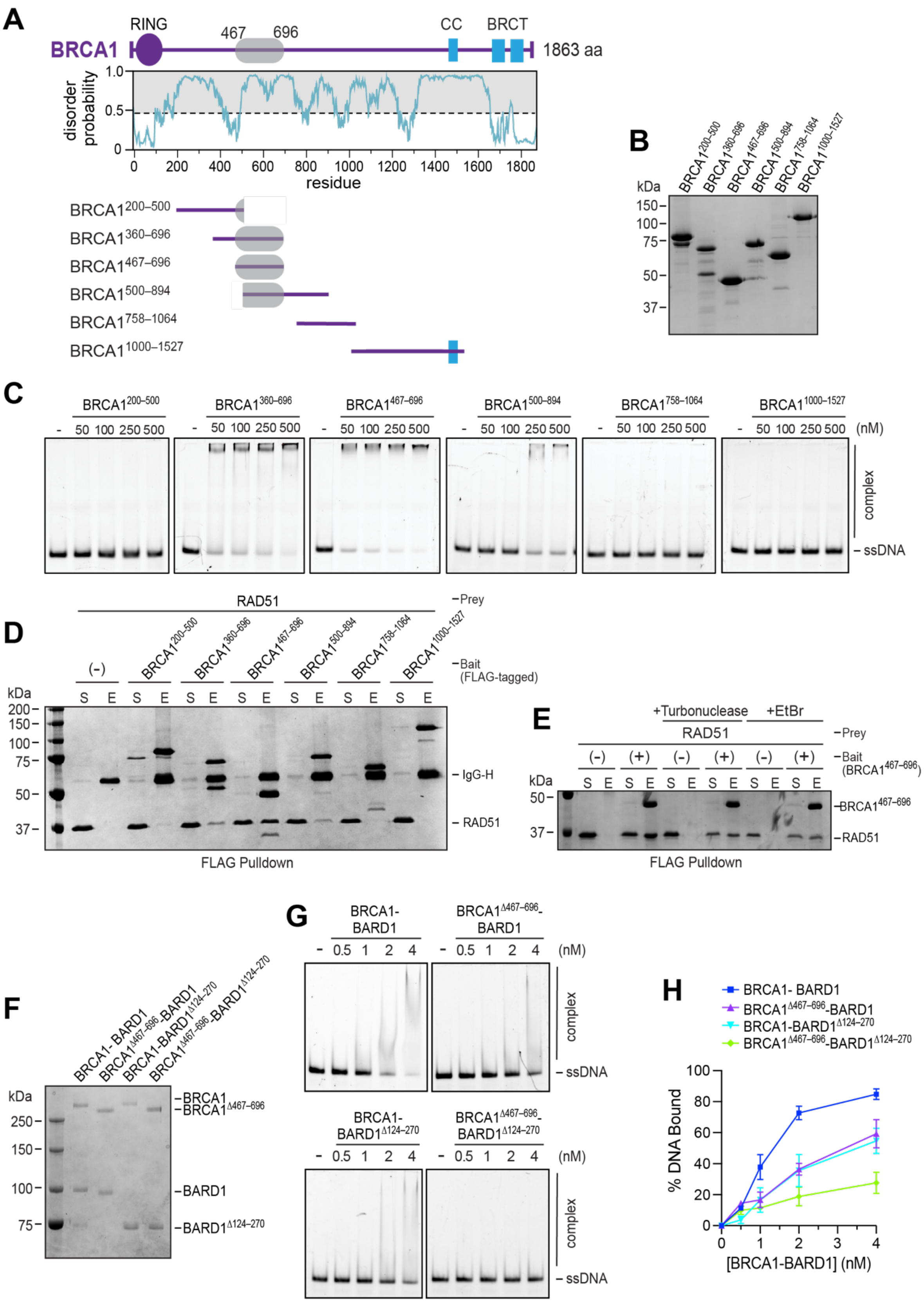

**Supplementary Figure 1. Identification of DNA binding and RAD51 interaction regions of BRCA1. Related to Figure 1.** (A) Disorder propensity of BRCA1 predicted using AIUPred[1] and schematic of the BRCA1 fragment library. (B) SDS-PAGE analysis of purified BRCA1 fragments, stained with Coomassie blue. (C) EMSA of purified BRCA1 fragments tested for DNA binding (Cy5-labeled 60-mer, 3 nM). (D) Testing of purified FLAG-tagged BRCA1 fragments tested for RAD51 interaction by affinity pulldown using anti-FLAG resin. Supernatant (S) and SDS eluate (E) were analyzed by SDS-PAGE with Coomassie blue staining. (E) Affinity pulldown testing interaction between RAD51 and $BRCA1^{467–696}$ in the presence of ethidium bromide or Turbonuclease, analyzed as in D. (F) SDS-PAGE analysis of BRCA1-BARD1 and mutants that harbor the indicated internal deletion. The gel was stained with Coomassie blue. (G) EMSA of BRCA1-BARD1 and the indicated DBR deletion mutants tested for dsDNA binding (Cy5-labeled 80-mer, 1 nM) by EMSA. (H) Quantification of results from G; n = 3; mean ± SEM.

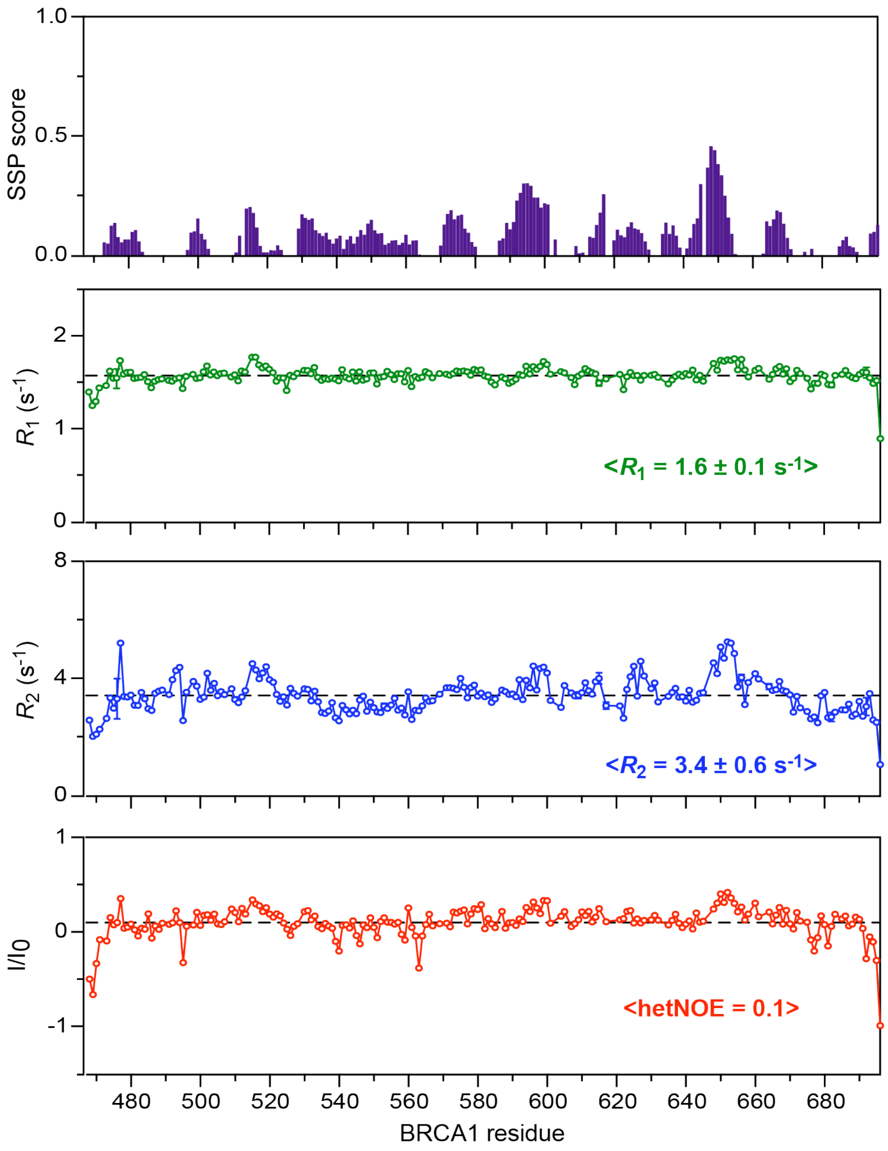


**Supplementary Figure 2. NMR characterization of $BRCA1^{467–696}$. Related to Figure 2.** Secondary structure propensity (SSP) of $BRCA1^{467–696}$ calculated from $^{13}C$ backbone chemical shifts. $^{15}N$ relaxation parameters including $R_1$ and $R_2$ rates and $^{1}H$-$^{15}N$ heteronuclear NOE (hetNOE) values plotted as a function of residue number.

### DNA binding-impaired mutants

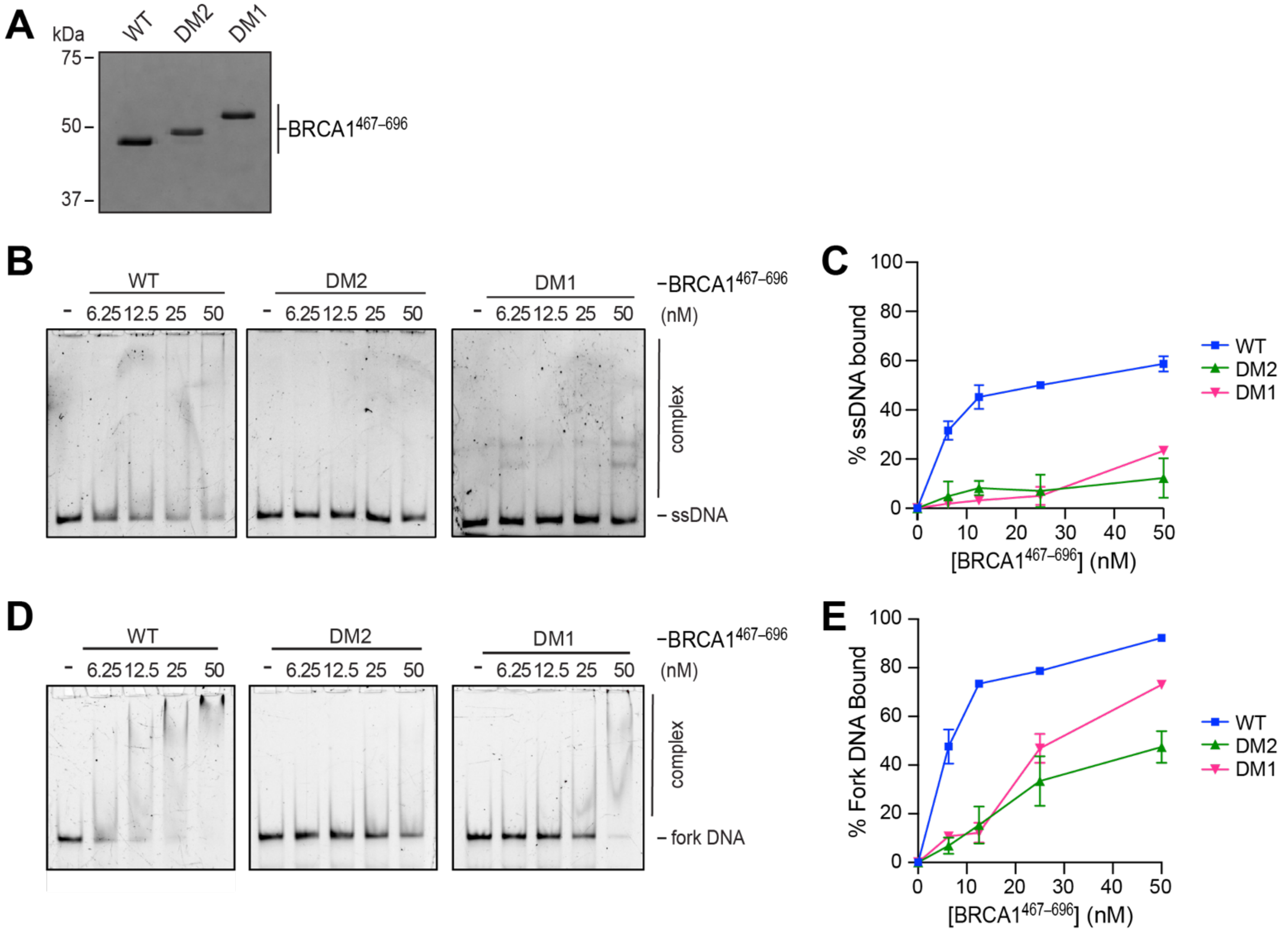


### RAD51 binding-impaired mutants

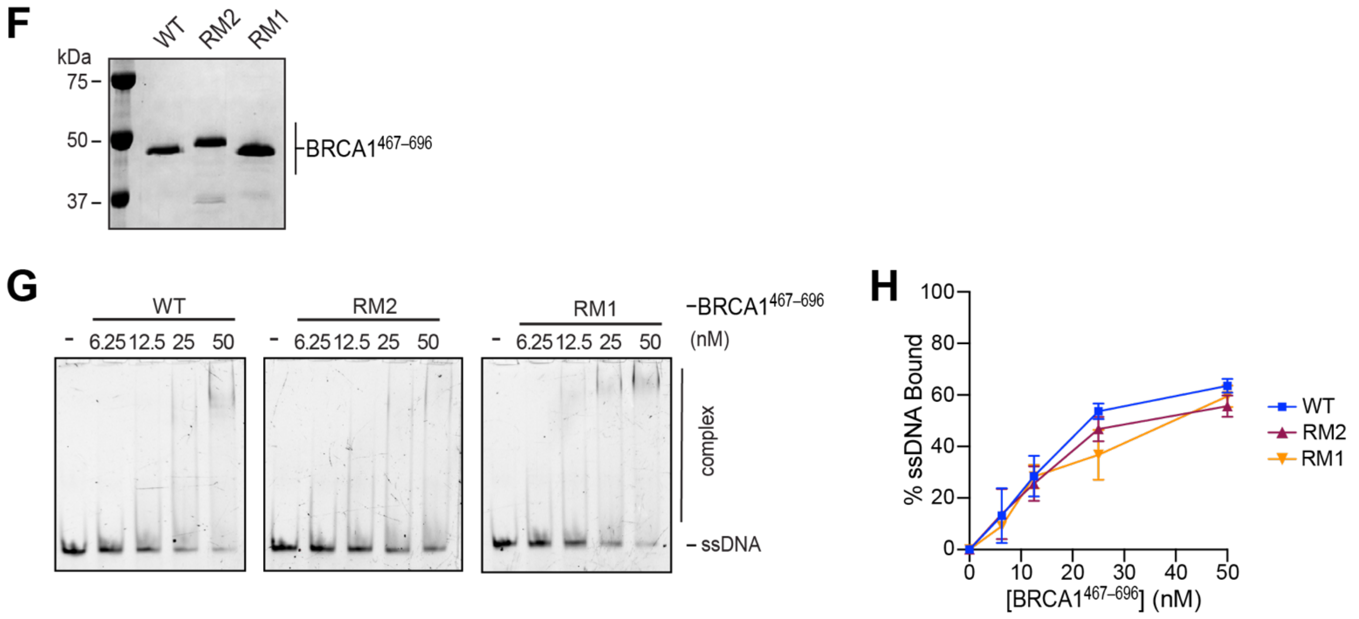

**Supplementary Figure 3. DNA binding characterization of BRCA1 separation-of-function mutants. Related to Figures 2, 3 and 4.** (A) SDS-PAGE analysis of wild type (WT), DM1, and DM2 variants of $BRCA1^{467–696}$ with Coomassie blue staining. (B) Testing $BRCA1^{467–696}$ WT, DM1, and DM2 for ssDNA binding (Cy5-labeled, 1 nM) by EMSA. (C). Quantification of results from B; n = 3; mean ± SEM. (D) $BRCA1^{467–696}$ WT, DM1, and DM2 tested for DNA fork binding (Cy5-labeled, 1 nM) by EMSA. (E) Quantification of results from D; n = 3; mean ± SEM. (F) SDS-PAGE analysis of $BRCA1^{467–696}$ WT, RM1, and RM2, stained with Coomassie blue. (G) Testing of $BRCA1^{467–696}$ WT, RM1, and RM2 for ssDNA binding (Cy5-labeled, 1 nM) by EMSA. (H) Quantification of results from G; n = 3; mean ± SEM.

### DNA binding-impaired mutants

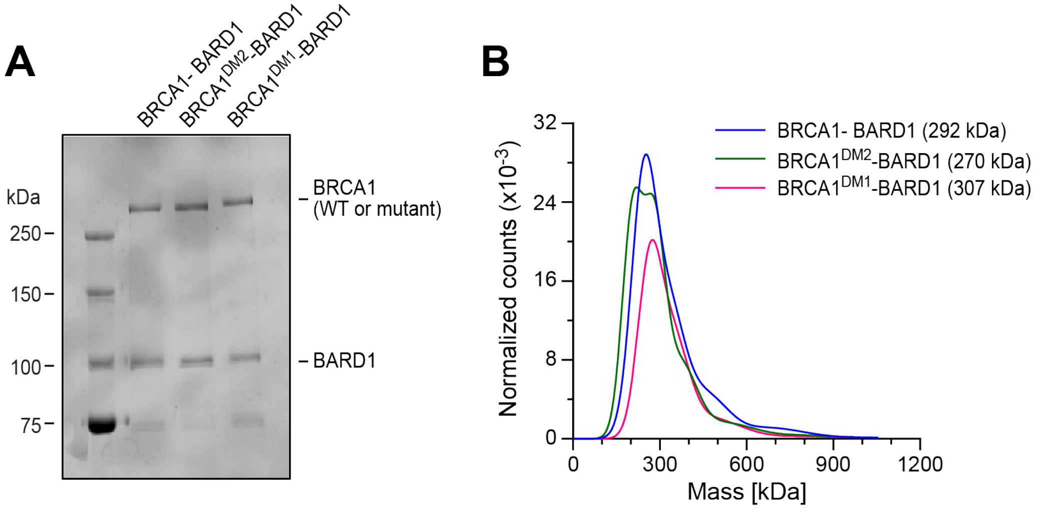


### RAD51 binding-impaired mutants

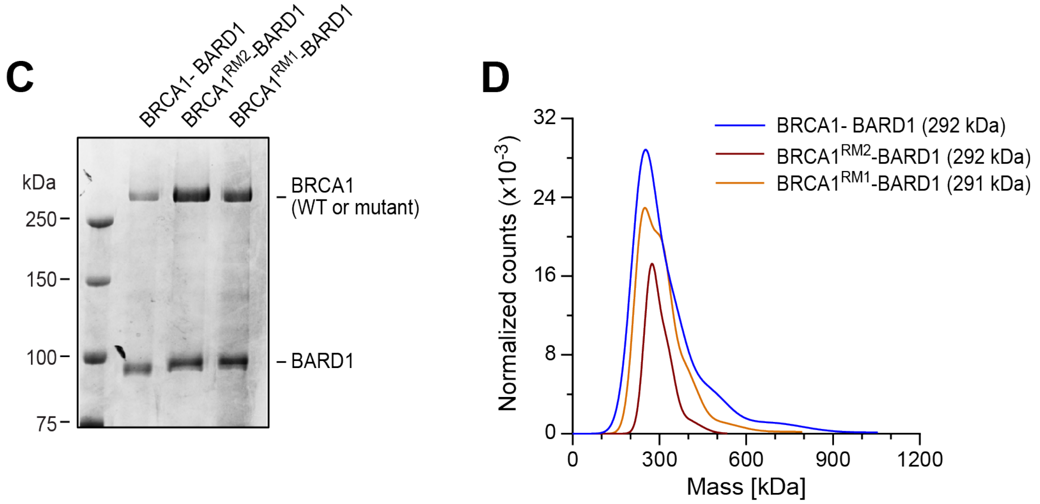


**Supplementary Figure 4. Analysis of full-length BRCA1-BARD1 SOF mutant complexes by mass photometry. Related to Figures 4 and 5.** (A) SDS-PAGE analysis of BRCA1-BARD1, BRCA1$^{DM1}$-BARD1, and BRCA1$^{DM2}$-BARD1, stained with Coomassie blue. (B) Mass photometry analysis of the complexes in A. (C) SDS-PAGE analysis of BRCA1-BARD1, BRCA1$^{RM1}$-BARD1, and BRCA1$^{RM2}$-BARD1, stained with Coomassie blue. (D) Mass photometry analysis of the complexes in C.

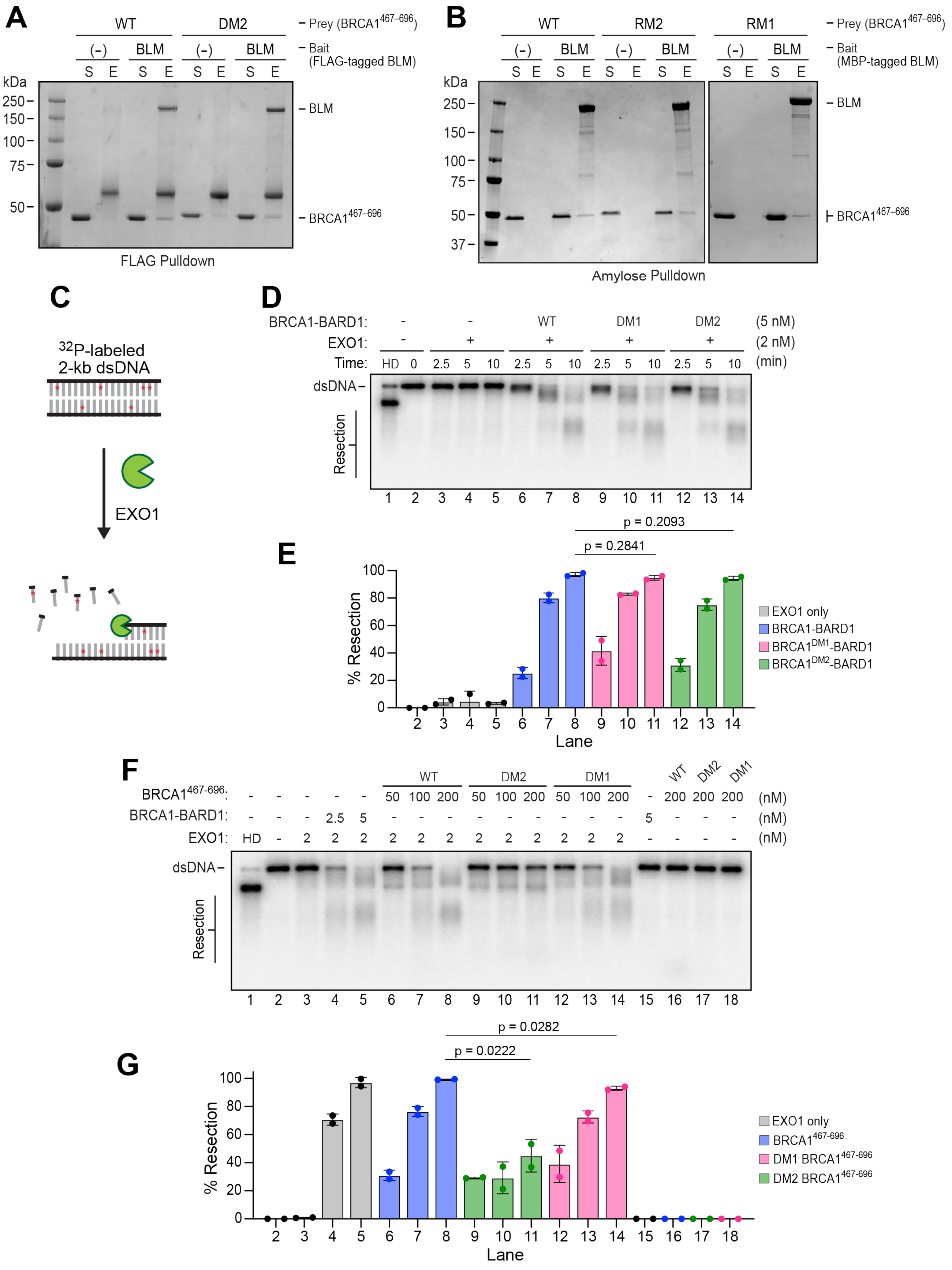

A
WT
DM2
(-)
BLM
S E S E S E S E
Prey (BRCA1467–696)
Bait (FLAG-tagged BLM)
kDa
250
150
100
75
50
BLM
BRCA1467–696
FLAG Pulldown
B
WT
RM2
RM1
Prey (BRCA1467–696)
Bait (MBP-tagged BLM)
kDa
250
150
100
75
50
37
BLM
BRCA1467–696
Amylose Pulldown
C
32P-labeled 2-kb dsDNA
EXO1
D
BRCA1-BARD1: - - WT DM1 DM2 (5 nM)
EXO1: - + + + + (2 nM)
Time: HD 0 2.5 5 10 2.5 5 10 2.5 5 10 2.5 5 10 (min)
dsDNA
Resection
1 2 3 4 5 6 7 8 9 10 11 12 13 14
E
p = 0.2093
p = 0.2841
% Resection
Lane
EXO1 only
BRCA1-BARD1
BRCA1DM1-BARD1
BRCA1DM2-BARD1
F
BRCA1467-696: - - - - - 50 100 200 50 100 200 50 100 200 - 200 200 200 (nM)
BRCA1-BARD1: - - - 2.5 5 - - - - - - - - - 5 - - - (nM)
EXO1: HD - 2 2 2 2 2 2 2 2 2 2 2 2 - - - - (nM)
WT DM2 DM1
dsDNA
Resection
1 2 3 4 5 6 7 8 9 10 11 12 13 14 15 16 17 18
G
p = 0.0282
p = 0.0222
% Resection
Lane
EXO1 only
BRCA1467-696
DM1 BRCA1467-696
DM2 BRCA1467-696

**Supplementary Figure 5. Role of BRCA1 DNA binding in DNA end resection. Related to Figure 5.** (A) Testing the interaction between wild-type (WT) and DM2 variants of $BRCA1^{467–696}$ with FLAG-tagged BLM using anti-FLAG M2 agarose. Supernatant (S) and SDS eluate (E) were analyzed by SDS-PAGE with Coomassie blue staining. (B) Testing the interaction between the WT, RM1, and RM2 variants of $BRCA1^{467–696}$ and MBP-tagged BLM using amylose resin, analyzed as in A. (C) Schematic of the EXO1 end resection assay. (D) BRCA1-BARD1, $BRCA1^{DM1}$-BARD1, and $BRCA1^{DM2}$-BARD1 tested for stimulation of EXO1-mediated end resection. (E) Quantification of results from D; n = 3; mean ± SEM. (F) $BRCA1^{467–696}$ WT, DM1, and DM2 tested for stimulation of EXO1-mediated end resection. (G) Quantification of results from F; n = 3; mean ± SEM.

### SUPPLEMENTARY MATERIAL REFERENCES


1. Erdos, G., and Dosztanyi, Z. (2024). AIUPred: combining energy estimation with deep learning for the enhanced prediction of protein disorder. Nucleic Acids Res *52*, W176–W181. 10.1093/nar/gkae385.
